# Single-neuron encoding of rapidly learned visual information reshapes human perception

**DOI:** 10.1101/2025.08.04.668333

**Authors:** Marcelo Armendariz, Julie Blumberg, Jed Singer, Franz Aiple, Jiye Kim, Nuria Dominguez-Iturza, Armin Brandt, Peter Reinacher, Andreas Schulze-Bonhage, Gabriel Kreiman

**Author notes:** These authors contributed equally (Marcelo Armendariz, Julie Blumberg).

## Abstract

Humans can swiftly learn to recognize visual objects after minimal exposure. Integrating new information with existing knowledge requires forming enduring neuronal representations to enable future recognition. Yet, the neuronal mechanisms in the human brain underlying such rapid perceptual changes remain unclear. We recorded single-neuron activity in occipital (OC) and medial temporal lobe (MTL) regions as participants rapidly learned to recognize degraded images. OC and MTL neurons modulated their activity to encode newly learned visual information and reshape perception. Population decoding revealed that OC neurons required additional processing time to resolve the identity of learned images, delaying neuronal responses in the MTL. Our findings indicate that OC supports recognition after rapid learning via extensive recurrent processing, potentially involving higher-order neocortical areas, without relying on feedback from MTL. These results provide mechanistic constraints for biologically plausible models of visual recognition and few-shot visual learning.

## INTRODUCTION

Humans and other primates display a remarkable ability to visually recognize objects. The main family of models to describe visual recognition capabilities is based on hierarchical deep neural networks^1–6^. Current instantiations of such networks are typically trained via backpropagation through millions of examples when learning from scratch but also when fine tuning an existing pre-trained network ^7–10^.

In stark contrast to such long training processes, humans can swiftly learn to recognize visual objects with just one or a few exposures. A striking example of rapid learning is the sudden recognition of a degraded black-and-white image of an object (Mooney image, **Figure 1a**)^11^. These degraded Mooney images tend to be unrecognizable initially. However, Mooney images become easily interpretable after a brief exposure to the original intact version of the image^12–14^ (**Figure 1b**). This perceptual ability for rapid learning develops gradually in late childhood (7-9 years) and is crucial for integrating new information with prior knowledge^15,16^. Rapid learning necessitates the formation of enduring neuronal signatures to enable subsequent recognition. Thus, newly acquired experience embedded in neural circuits dramatically shapes the perception of future incoming visual stimuli^15,17–19^. Despite extensive behavioral characterization, the neuronal mechanisms underlying perceptual changes induced by rapid learning in the human brain are not well understood.

**Figure 1.**
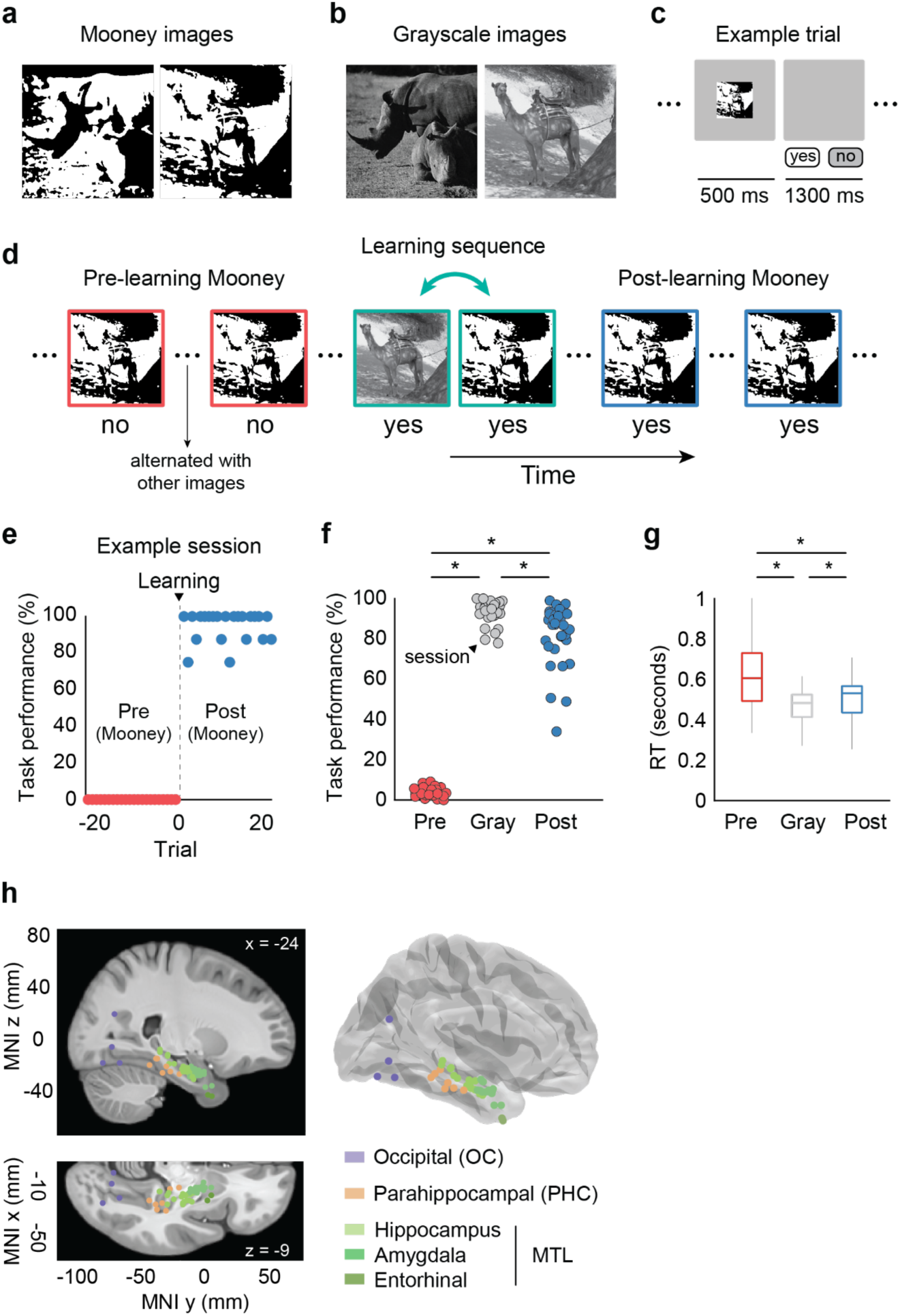
Experimental design, behavior, and recording sites. **a**, Examples of Mooney images. **b**, Examples of Grayscale images. **c**, Example trial displaying stimulus and blank screen durations. Participants reported whether they recognized the object on the screen via button press (’yes’ or ‘no’). **d**, Illustration of the learning process for Mooney images. The sequence depicts Mooney images before learning (Pre-learning, red frame) and after learning (Post-learning, blue frame), with the learning phase marked by a turquoise double-headed arrow. Pre-learning Mooney images are unrecognizable, as indicated by the response ‘no’ shown beneath the image. After multiple presentations of Pre-learning Mooney images, Grayscale and a Mooney image depicting the same object are shown back-to-back to facilitate learning of the Mooney image. During the learning sequence, both the Grayscale and Mooney images are recognized (’yes’ beneath the images). Following the learning sequence, Mooney images can be recognized. The colored frames were added for illustration purposes and were not visible during the task. Note that this panel illustrates a sequence for a single image (camel); the full sequence included multiple images interleaved throughout the session, as indicated by the ellipses between stimuli (see Methods). **e**, Proportion of Mooney images recognized across trials during an example session. Trials are aligned with respect to the learning event (trial 0), indicated by a downward arrow. Mooney image recognition rapidly and drastically grew from almost none to almost perfect after learning. Post-learning trials (positive numbers, blue dots) show proportion of Mooney images recognized after learning. Pre-learning trials (negative numbers, red dots) show that later-recognized Mooney images were not recognized before learning, even though the images were identical. **f**, Behavioral performance during the recognition task across all sessions. Before learning (Pre, red), participants exhibited low performance (4%). Grayscale images (Gray, gray) and Mooney images after learning (Post, blue) were highly recognizable (93% and 83%, respectively). Each dot represents a session (N=34). **g**, Response times (RT) relative to image onset. Participants responded faster to Post-learning Mooney images compared to Pre-learning Mooney images, and the fastest responses were those to Grayscale images. Box plots show the median and interquartile range of the response time distributions. Asterisks (*) in **f** and **g** show statistically significant differences between conditions (*P* < 0.05, two-sided Wilcoxon signed-rank test). **h**, Electrode locations across all participants. Each dot represents a microwire bundle location, mapped onto a standardized MNI152 brain template (left) and a 3D brain model (right). All dots are displayed in the same hemisphere for visualization purposes only. The color of each dot denotes the microwire bundle location.

Previous neurophysiological recordings in monkeys showed that neurons in the temporal visual cortex can adapt their selectivity to represent recently learned degraded versions of objects such as Mooney images^20^. Based on non-invasive methods, rapid learning in humans is thought to modulate activity in a broad range of neocortical regions, including visual areas and frontoparietal regions^21–24^. However, little is known about the mechanisms underlying how individual neurons modify their response patterns after learning and how rapid and all-or-none perceptual changes are orchestrated by the neuronal dynamics across different regions.

Here we recorded the spiking activity of 1,104 single neurons in medial occipital and temporal regions of the human brain in patients with pharmacologically-resistant epilepsy while 13 participants learned to rapidly recognize novel Mooney images. Participants demonstrated the hallmark characteristics of essentially all-or-none perceptual changes triggered by one or a few exposures to grayscale counterparts to the Mooney images. Neurons in both the OC and MTL modulated their responses to encode newly learned Mooney images. OC neurons required additional time to process the identity of learned Mooney images compared to intact grayscale images, with MTL neurons showing delayed responses. The learning-induced dynamics in OC may indicate the need for extensive recurrent processing, potentially involving top-down feedback from higher cortical areas, before information reaches MTL.

## RESULTS

### Rapid learning during image recognition and single neuron recordings

We designed an image recognition task to investigate changes in perception following rapid learning. Thirteen participants were shown sequences of images and were instructed to report whether they recognized the identity of the depicted objects. Images consisted of two-tone black and white pictures (Mooney images, **Figure 1a**) and their corresponding grayscale counterparts (**Figure 1b**). In each trial, an image was shown for 500 milliseconds, followed by a blank screen for 1,300 milliseconds (**Figure 1c**). Participants were first exposed to the Mooney versions of the images, which they did not recognize (**Figure 1d**, red). After exposure to the Mooney images, brief learning segments were introduced. These segments involved presenting participants with the original grayscale image followed by the corresponding Mooney version of that image (**Figure 1d**, turquoise). After exposure to these learning segments, participants were shown the same Mooney images again (**Figure 1d**, blue). Additional grayscale images were also presented, though no longer back-to-back with the corresponding Mooney images. Different Mooney and grayscale images were interleaved throughout the task. Using this design, we defined three trial categories: Pre-learning, where Mooney images were presented *before* learning occurred; Grayscale, where the original grayscale images were presented; and Post-learning, where Mooney images were presented *after* the learning process. Mooney image recognition rapidly and drastically grew from almost none before learning to high levels after learning (**Figure 1e**), requiring only one or a few learning segments (on average, 1.8 ± 1). Overall, Grayscale images were highly recognizable, with participants performing on average at 93 ± 5% accuracy (mean ± s.d. across sessions, n = 34 sessions; **Figure 1f**). For the Mooney images, recognition was poor in the Pre-learning condition with an accuracy of 4 ± 2% and greatly improved for the Post-learning condition, reaching 83 ± 15%. Performance was statistically different between conditions (Gray vs Pre, *P* = 10^-12^; Gray vs Post, *P* = 10^-4^; Pre vs Post, *P* = 10^-12^; two-sided Wilcoxon signed-rank test). Additionally, we measured reaction times from stimulus onset to participants’ responses (**Figure 1g**). Response times were longer for recognized Mooney images (Post-learning: median = 0.53 s, IQR = 0.44–0.57 s; mean = 0.52 s) than for Grayscale images (median = 0.48 s, IQR = 0.41–0.52 s; mean = 0.48 s) (Post vs Gray, *P* = 10^-5^; two-sided Wilcoxon signed-rank test), suggesting that interpreting the learned Mooney stimuli required additional cognitive effort. Unrecognized Mooney images (Pre-learning) elicited the slowest responses overall (median = 0.60 s, IQR = 0.49–0.72 s; mean = 0.63 s) (Pre vs Gray, *P* = 10^-7^; Pre vs Post, *P* = 10^-7^ two-sided Wilcoxon signed-rank test).

During this task, we recorded spiking activity from a total of 1,104 single neurons in medial occipital and medial temporal brain areas in 34 sessions from 13 patients with pharmacologically-resistant epilepsy who were chronically implanted with hybrid macro- and microelectrodes for seizure monitoring (**Figure 1h; Supplementary Table 1**). The recordings included 163 neurons from the medial occipital cortex, 212 in the parahippocampal cortex, 49 in the entorhinal cortex, 343 in the hippocampus, and 337 in the amygdala (**Supplementary Table 2**).

We characterized eye movements during the task in 13 sessions across 10 healthy participants. These participants exhibited similar recognition patterns to the 13 epilepsy patients, with high recognition of Mooney images only after learning, and longer reaction times for recognized Mooney images compared to Grayscale images (compare **Supplementary Figure 1** and **Figure 1**). In the majority of trials (∼83%), participants did not make any saccades during image presentation (0–500 ms). In the subset of trials with saccades (∼17%), the latency to the first saccade from image onset was 267 ± 33 ms. There were no significant differences across conditions in either the proportion of trials with saccades (Gray vs Pre, *P* = 0.17; Gray vs Post, *P* = 0.52; Pre vs Post, *P* = 0.38; two-sided Wilcoxon signed-rank test) or the timing of the first saccade (Gray vs Pre, *P* = 0.49; Gray vs Post, *P* = 0.99; Pre vs Post, *P* = 0.49; two-sided Wilcoxon signed-rank test).

### Neuronal latency and selectivity reflect hierarchical processing across brain regions

We first identified visually responsive neurons. A neuron was considered responsive if its firing rate showed significant modulation to any of the presented images, during the interval from 200 to 700 ms after image onset, compared to the baseline period (−500 to 0 ms before image onset). Significance was determined using a permutation test (1,000 iterations) with a threshold P-value < 0.05, corrected for false discovery rate (FDR). We identified 318 visually responsive neurons (29% of the total). Of these, 105 were located in the medial occipital cortex (OC, 64% of the total number of neurons in this area), 90 in the parahippocampal cortex (PHC, 42%), 53 in the hippocampus (15%), 59 in the amygdala (17%), and 11 in the entorhinal cortex (22%) (**Figure 2a**). We will refer to the neurons in the hippocampus, amygdala, and entorhinal cortex together as the medial temporal lobe (MTL), comprising 123 visually responsive neurons. Representative examples are shown in **Supplementary Figure 2a-d**. For instance, **Supplementary Figure 2a** illustrates the responses of a hippocampal neuron to a specific image in its Pre Mooney, Grayscale, and Post Mooney versions. This neuron exhibited distinct response patterns across conditions, with highest responses observed in the Grayscale condition. Subsequent analyses were based only on visually responsive neurons.

**Figure 2.**
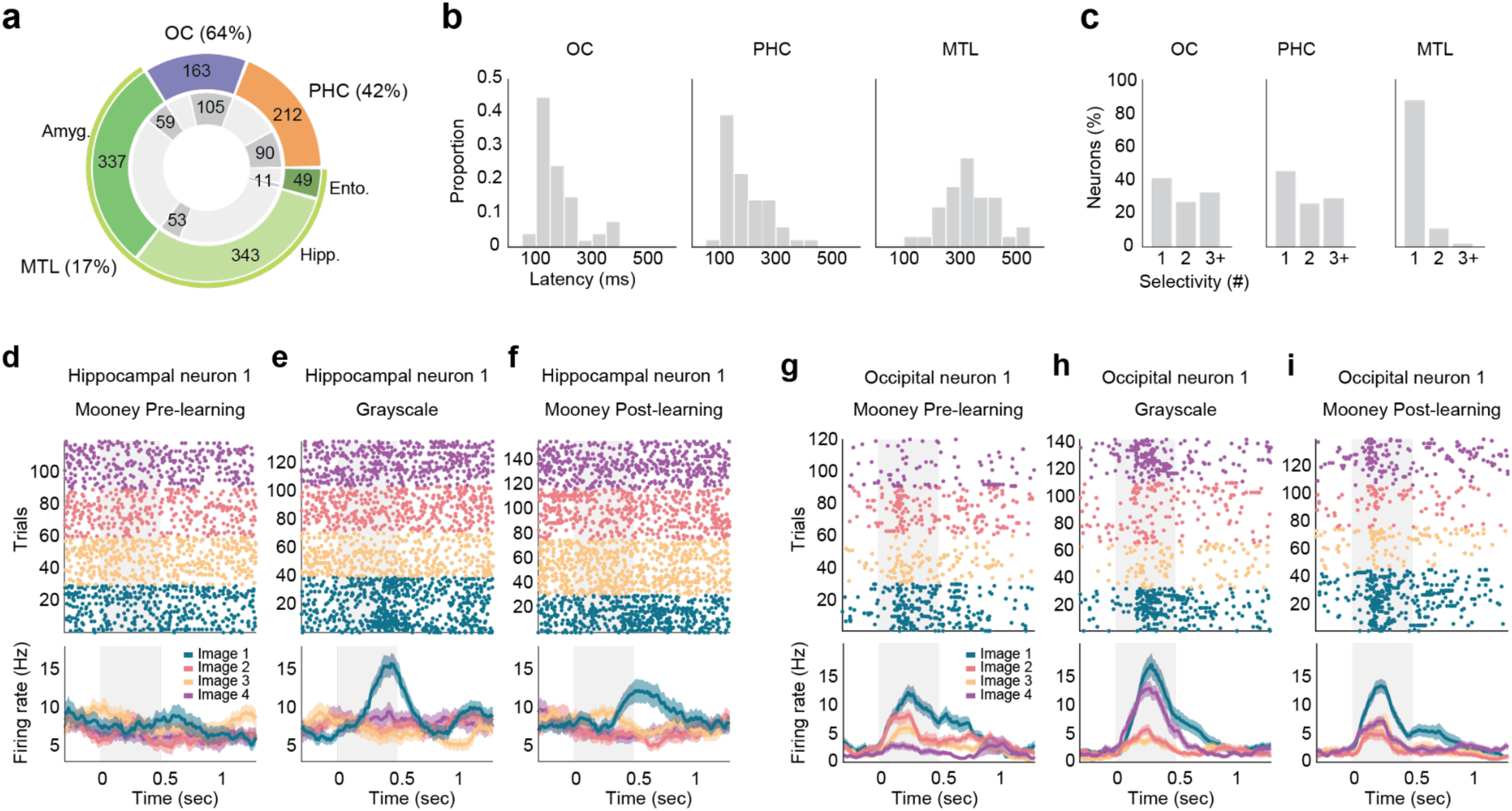
Neurons show different response latencies and visual selectivity across brain regions. **a**, Proportion (and count) of visually responsive neurons across human brain regions. OC refers to the occipital cortex, and PHC to the parahippocampal cortex. Neurons in the hippocampus (Hipp.), amygdala (Amyg.), and entorhinal cortex (Ento.) are collectively referred to as the MTL from here on. Visually responsive neurons showed statistically significant modulation of their firing rates during image presentation (Grayscale or Mooney) compared to the baseline period (*P* < 0.05, FDR corrected, permutation test, 1,000 iterations). The ring chart shows the number of visually responsive neurons (dark gray segments) relative to the total number of recorded neurons (colored segments). Percentages indicate the proportion of responsive neurons within each brain region (reported in aggregate for the MTL). **b**, Distribution of neuronal response latencies across brain regions. Response latency was defined as the time from stimulus onset to the first significant change in firing rate relative to baseline (Methods). **c**, Proportion of neurons across brain regions that selectively responded to 1, 2 or 3+ distinct grayscale images. **d**,**e**,**f**, Example of a highly selective neuron in the hippocampus. Panels show the spiking activity in response to different images (Images 1-4) in the Pre-learning Mooney (**d**), Grayscale (**e**), and Post-learning Mooney (**f**) conditions. This neuron responded only to Image 1 (turquoise) for the Grayscale (**e**) and Post-learning Mooney (**f**) images, but not in the Pre-learning condition (**d**). Raster plots (top) show spiking activity (each dot represents a spike event) across trials (rows) and over time. Colors correspond to different images. Responses are shown for a subset of four images that were presented during the task. Trials are grouped by image for visualization. The light gray rectangle in the background indicates the image presentation interval. Mean firing rates (bottom) are shown separately for each image (solid colored lines; bin size is 250 ms and step size is 10 ms). Shaded areas represent s.e.m. across trials. Firing rates for the Mooney images increased after learning (**f**) compared to before learning (**d**). **g**,**h**,**i**, Spiking activity of an example neuron in the medial occipital cortex, shown with the same conventions as in **d**-**f**. This neuron responded to multiple stimuli (Images 1-4) and exhibited firing rate modulation depending on the image and condition.

We next estimated the onset latencies for responsive neurons. Response latency was defined as the first significant change in the neuron’s firing rate relative to the baseline period after stimulus onset for the grayscale images (Methods). Response latencies significantly differed between areas with the fastest responses in the OC (median = 155 ms, interquartile range IQR = 129–204 ms), followed by the PHC (median = 161 ms, IQR = 138–248 ms), and the MTL (median = 322 ms, IQR = 272–392 ms) (*P* = 10^-10^, Kruskal–Wallis test, **Figure 2b**).

Next, we evaluated the degree of visual selectivity of responsive neurons. We assessed response selectivity based on the number of distinct grayscale images a neuron responded to. MTL neurons showed the highest degree of selectivity with 87% of responsive neurons selective to only one stimulus, followed by neurons in PHC (45%) and OC (40%) (**Figure 2c**). **Figure 2d-i** shows the spiking activity (top, raster plots; bottom, mean firing rates) of two example neurons in response to different images (color-coded, Images 1-4) and across conditions. **Figure 2d-f** illustrates a highly selective neuron in the hippocampus. In the Grayscale condition, this neuron responded only to Image 1 (turquoise, **Figure 2e**). A second example, **Figure 2g-i**, shows the spiking activity of a neuron in the medial occipital cortex that responded to multiple images. This neuron exhibited firing rate modulation depending on the image and condition. For instance, responses to images 1 (turquoise) and 4 (purple) were higher in the Grayscale condition (**Figure 2h**) compared to the Mooney conditions (**Figure 2g,i**). However, for image 3 (yellow) firing rates were slightly lower in the Grayscale condition compared to the Mooney conditions. Additional example neurons are shown in **Supplementary Figures 2 and 3**. Overall, these results suggest a hierarchical organization in visual processing, where both selectivity and latencies increase as neural signals flow from the occipital region to the medial temporal lobe.

### Neuronal activity is modulated by learning at the single neuron level and in single trials

Modulation in the responses to grayscale images compared to their Mooney counterparts (**Figure 2e** vs. **Figure 2d,2f**; **Figure 2h** vs. **Figure 2g,i**; **Supplementary Figure 3b** vs. **Supplementary Figure 3a,c**; **Supplementary Figure 3e** vs. **Supplementary Figure 3d,f**) could be expected and may reflect the large differences between the two types of images at the pixel level. Remarkably, these differences were also accompanied by modulation in firing rates between the identical Post- and Pre-learning Mooney images. The hippocampal neuron in Figure 2 remained unresponsive to all Mooney images *before* learning (**Figure 2d**), but its firing rate was selectively modulated in response to Mooney Image 1 *after* learning (**Figure 2f**). Similarly, the occipital neuron in **Figure 2** did not respond to Mooney Image 4 (purple) before learning (**Figure 2g**) but increased its firing rate in response to this Mooney image after learning (**Figure 2i**). See **Supplementary Figure 2** for four examples showing a direct comparison between the responses to specific images across conditions and **Supplementary Figure 3** for further examples in the same format of **Figure 2**.

We systematically examined whether neurons modulated their activity after learning compared to before learning, as indicated by differences in firing profiles between Pre- and Post-learning Mooney conditions. Both conditions utilized the same stimuli, providing identical visual input. However, the latter occurred after learning and thus the depicted image identity could be readily recognized while the former elicited no recognition (**Figure 1d,e,f**). We hypothesized that these perceptual differences would be reflected in changes of firing patterns of individual neurons. To test this hypothesis, we first trained linear decoders (support vector machine with linear kernel, bin size = 250 ms and step size = 25 ms; Methods) separately for each neuron to discriminate between response patterns for Pre- and Post-learning trials across time (**Figure 3a**). On average, the single-neuron classifier could decode between the two conditions (*P* = 0.001, permutation test), evidencing the modulating effects of learning at the single neuron level and in single trials during image recognition.

**Figure 3.**
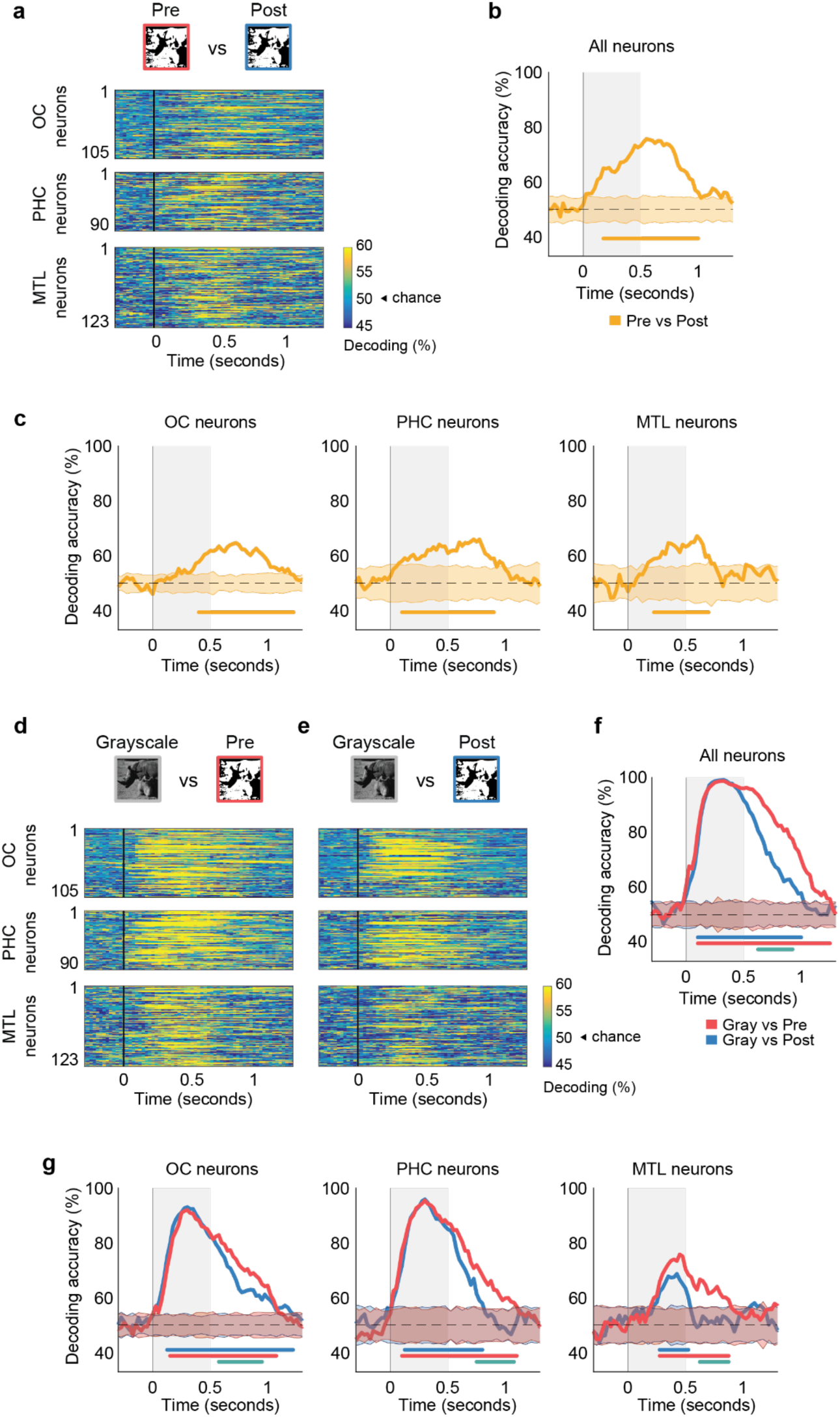
Neuronal activity is modulated by learning. **a**, Decoding accuracy for Pre- (red frame) versus Post-learning Mooney (blue frame) images. Each row corresponds to the decoding accuracy of the responses of single neurons in the medial occipital cortex (OC, top), parahippocampal cortex (PHC, middle) and the medial temporal lobe (MTL, bottom). See color scale on bottom right. **b**, Population decoding accuracy including the responses of all neurons in **a** as features. The dashed line is chance (50%). The solid line below the curves represents time points significantly above chance (*P* < 0.05, versus empirical null distribution). In this and following panels, time zero indicates image presentation onset and the shaded gray area represents the stimulus presentation period. **c**, Population decoding accuracy using neurons in **a** as features for each brain region separately. **d**, Decoding accuracy for Grayscale (gray frame) vs. Pre-learning Mooney images and **e**, Grayscale vs. Post-learning Mooney images. Same conventions and color scale as in **a**. **f**, Population decoding accuracy including the responses from all neurons in **d** (Gray vs Pre, red) and **e** (Gray vs Post, blue) as features. Same conventions as in **b**. The solid turquoise line below the curves represents the time points with statistically significant difference between the two decoding profiles (P < 0.05, versus empirical null distribution). **g**, Population decoding accuracy using the responses of neurons in **d** as features for each brain region separately. Same conventions as in **f**. In **b**, **c**, **f**, and **g**, the shaded area around the chance line represents the 95% confidence interval of the empirical null distribution obtained by shuffling the labels (500 iterations). Decoders were binary linear classifiers (support vector machine; chance = 50%) using sliding windows of 250 ms and a step size of 25 ms.

We next assessed the decoding ability of pseudopopulations consisting of groups of single neurons across all participants^25,26^ (Methods). We pooled all responsive neurons to create a pseudopopulation and built linear decoders using the responses of single neurons as features (bin size = 250ms, step size = 25ms; Methods). The classifier discriminated between Pre- and Post-learning images with an accuracy of up to 75% at 550 ms (**Figure 3b**). Reliable decoding started during image presentation (175-500ms) and continued after stimulus offset (500-900ms) (*P* < 0.05, permutation test versus empirical null distribution, chance = 50%).

Next, we investigated whether learning-induced modulation of neuronal activity was localized to specific brain regions. Neuronal pseudopopulations in the OC, PHC, and MTL could discriminate between Mooney images before versus after learning (**Figure 3c**). Decoding started during image presentation (*P* < 0.05, permutation test versus empirical null distribution), and peaked briefly after stimulus offset in the three regions. Taken together, these results show that newly acquired experience induces changes in neuronal activity, evident even in single trials, across brain areas spanning the human medial occipital and temporal lobes, commencing during stimulus presentation and continuing beyond the stimulus offset.

### Learning aligns neuronal patterns for Mooney images to Grayscale images

We further investigated the nature of learning-induced firing rate modulation by comparing neuronal response patterns for Pre- and Post-learning Mooney images with those of the Grayscale images. Given the distinct stimuli for Mooney images compared to Grayscale images, one might expect dissimilar neuronal patterns purely due to visual input differences. On the other hand, since participants could recognize both Grayscale and Post-learning Mooney images but not Pre-learning ones (**Figure 1f**), we hypothesized that learning-induced response modulation might align firing rates of Post-learning Mooney images with those of Grayscale images to support recognition. Such alignment would result in *less* discriminability between neuronal patterns for Post-learning and Grayscale images compared to discriminability between Pre-learning and Grayscale images. Examination of the example neurons in **Figure 2, Supplementary Figure 2**, and **Supplementary Figure 3**, is consistent with this idea. The hippocampal neuron in **Figure 2d-f** responded only to Image 1 both for the Grayscale (**Figure 2e**) and Post-learning Mooney images (**Figure 2f**), but not for the Pre-learning Mooney image (**Figure 2d**). The occipital neuron in **Figure 2g-i** responded to Image 4 only for the Grayscale (**Figure 2h**) and Post-learning Mooney images (**Figure 2i**).

To test the hypothesis of response alignment across all neurons, we first trained linear decoders at the single neuron level to discriminate Grayscale images from Mooney images, separately for Pre- and Post-learning conditions. On average, neurons could reliably discriminate between Grayscale and Mooney images in both Pre- (**Figure 3d**) and Post-learning (**Figure 3e**) conditions (*P* = 0.001, permutation test). Consistent with the hypothesis, average decoding accuracy was overall higher for the Pre-learning condition compared to the Post-learning condition, both during image presentation and after image offset (**Supplementary Figure 4**). A pseudopopulation analysis using all neurons as features reinforced these findings (**Figure 3f**). Neuronal response patterns between Mooney and Grayscale images were rapidly (125 ms) and highly decodable (98%, *P* < 0.05, permutation test versus empirical null distribution), and lasted longer for the Pre-learning condition compared to the Post-learning condition. The dynamics of decoding profiles were similar for Pre- and Post-learning conditions during image presentation. The difference between Pre- and Post-learning conditions, evident in **Supplementary Figure 4** during stimulus presentation, may be obscured in **Figure 3f** due to an accuracy ceiling effect. Decoding accuracies at the pseudopopulation level began to diverge shortly after the stimulus disappeared from the screen, exhibiting a significant decrease in discriminability for the Post-learning condition relative to the Pre-learning condition (*P* < 0.05, permutation test versus empirical null distribution). In line with the proposed hypothesis, the drop in decoding for the Post-learning Mooney images indicates that neuronal response patterns evoked by the Post-learning images tended to align to those evoked by Grayscale images.

We repeated these analyses separately for each brain region (**Figure 3g**). OC and PHC showed high discriminability between Grayscale and both Pre- and Post-learning Mooney conditions during the image presentation phase, with divergent decoding profiles after stimulus offset (*P* < 0.05, permutation test versus empirical null distribution). In the MTL, the Gray-vs-Pre and Gray- vs-Post decoding profiles began to diverge consistently earlier during image presentation, but significant differences only emerged after image offset. For both profiles, discriminability between Grayscale and Mooney images was observed earlier in the OC and PHC (125-175 ms) compared to the MTL (300ms). Overall, following learning, neurons changed their firing patterns in response to Mooney images to resemble those of the Grayscale images across all three brain regions.

### Neuronal patterns for Grayscale images generalize to Mooney images after learning

To further understand the content encoded by neuronal populations before versus after learning, we assessed whether activity patterns elicited by Grayscale images generalized to two-tone Mooney images. If neurons changed their firing patterns after learning to reflect image identity, then patterns encoding Grayscale images would be expected to generalize to Mooney images after learning, but not (or to a lesser extent) before learning. To test this hypothesis, we employed a cross-condition decoding approach in which classifiers were trained on one condition and tested on another across time (**Figure 4a-f**). At each time point and for every pair of conditions, decoders predicted image identity from neuronal activity patterns. Because different sessions used distinct subsets of images, we chose the top three images from each session based on their responses in the Grayscale condition and labeled the selected images of each session as Image 1-3 to build three-class decoders (Methods; chance = 33%).

**Figure 4.**
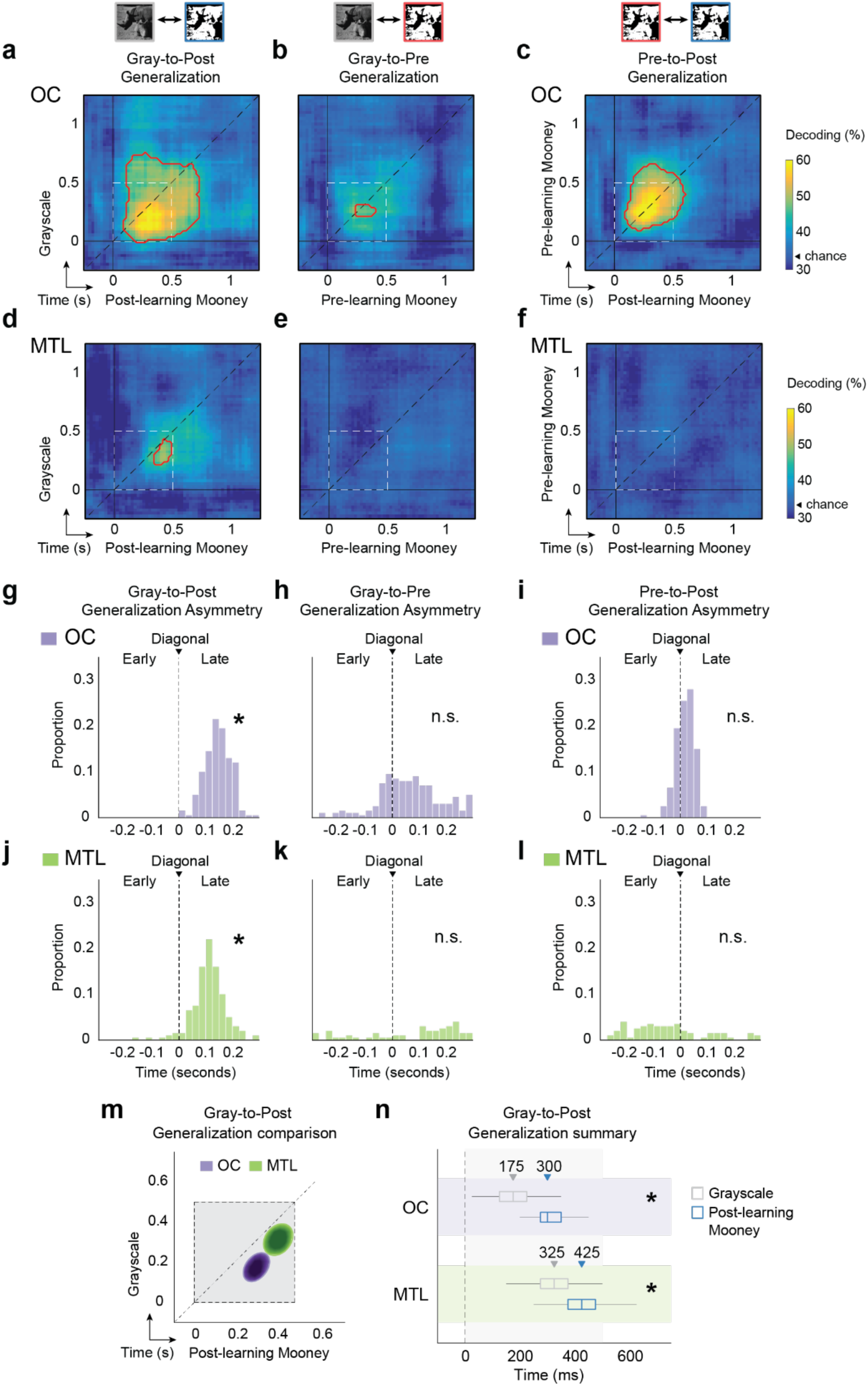
Generalization to learned Mooney images is delayed in OC and MTL. **a-c**, Generalization matrices in OC for: (**a)** Grayscale to Post-learning Mooney (Gray-to-Post), (**b**) Grayscale to Pre-learning Mooney (Gray-to-Pre), and (**c**) Pre-learning to Post-learning Mooney (Pre-to-Post). Generalization was assessed by training a decoder with one condition and testing with the other, including the responses of neurons as features. Each matrix entry *x,y* indicates the mean decoding accuracy of image identity for the two train-test configurations across conditions, as follows: first, we computed decoding accuracy by training with the Grayscale at time *y* and testing with the Mooney condition at time *x*; second, we computed decoding accuracy by training with the Mooney at time *x* and testing with the Grayscale condition at time *y*; third, the mean of these two values correspond to entry *x,y*. Accuracies correspond to three-class decoders (chance is 33%) of top three images chosen based on their responses in the Grayscale condition (Methods). The x- and y-axes indicate time points of neuronal activity for the corresponding conditions. Time zero marks image onset. The small pictures above the matrices indicate the conditions included in the generalization matrix (gray frame for Grayscale, blue frame for Post-learning Mooney, and red frame for Pre-learning Mooney). The black dashed line represents the matrix diagonal, where conditions were trained and tested at the same time points. The dashed line square represents the image presentation interval. Red solid lines delineate statistically significant decoding (*P* < 0.05, permutation test versus empirical null distribution). **d**,**e**,**f**, Generalization in MTL. Panels and data follow the same conventions as in **a**-**c**. **g**,**h**,**i**, Generalization asymmetry corresponding to decoding profiles in **a**, **b**, and **c**. We quantified the deviation of peak decoding from the diagonal to measure generalization asymmetry. Deviation of the distribution to the right indicates that neuronal patterns for Grayscale images generalize to later activity for the Post-learning Mooney images. Asterisks (*) indicate statistically significant deviation from the diagonal (*P* < 0.05, empirical distribution versus the diagonal). n.s indicates no statistically significant deviation from the diagonal. **j**,**k**,**l**, Generalization asymmetry in MTL corresponding to the decoding profiles in **d**, **e**, and **f**, respectively. Data use the same conventions as in **g**-**i**. Results for the OC are color-coded in purple, and MTL in green. **m**, Comparison of Gray-to-Post generalization in the OC and MTL. The ellipses illustrate generalization asymmetry in the OC (purple) and MTL (green), corresponding to panels **a** and **d**, respectively. In both cases, peak generalization falls below the diagonal, occurring earlier in the OC than in the MTL. The ellipses span the interquartile range of the probability density functions of peak generalization (**Supplementary Figure 6a,d**). The shaded gray area with dashed lines delineates the stimulus presentation period. **n**, Summary of generalization times for Gray-to-Post. Downward arrows and numbers indicate the median peak generalization times for Grayscale (gray box) and Post-learning Mooney (blue box) images in the OC (top, purple background) and MTL (bottom, green background). Box plots represent the interquartile range of the marginals of the probability density functions (**Supplementary Figure 6a,d**). The vertical dashed line at time zero indicates image presentation onset, and the shaded gray area in the background represents the presentation period.

We examined generalization of neuronal responses for Grayscale to Mooney images (Gray-to-Post and Gray-to-Pre) separately for each region. **Figure 4a** shows Gray-to-Post generalization for neurons in the OC. Each entry at time *x*, *y* indicates the mean decoding accuracy for two train-test configurations: first, the classifier was trained with the responses to Grayscale images at time *y* and tested with the responses to Post-learning Mooney images at time *x*; second, the classifier was trained with the responses to Post-learning Mooney images at time *x* and tested with the responses to Grayscale images at time *y*. OC neurons showed significant Gray-to-Post generalization (*P* < 0.05, versus empirical null distribution; chance = 33%; **Figure 4a**), and weaker generalization for Gray-to-Pre (**Figure 4b**). Decoding accuracies were higher for Gray-to-Post (up to 62%) than for Gray-to-Pre (49%), and remained significant for ∼200 ms after image offset for the Post-learning Mooney images. We next evaluated generalization in MTL neurons, which showed significant transfer of image identity information only after learning, with decoding accuracy reaching 50% for Gray-to-Post (*P* < 0.05; **Figure 4d**), and chance-level performance for Gray-to-Pre (**Figure 4e**). Finally, generalization analyses in the parahippocampal cortex revealed no significant decoding across conditions (**Supplementary Figure 5**).

Additionally, we asked whether neural responses would generalize from Pre- to Post-learning Mooney images, given that the visual input was identical despite the change in perception, from unrecognized to recognized. We found that Pre-to-Post generalization was significant in OC neurons (up to 65%; **Figure 4c**), with decoding mostly confined to the image presentation interval. In contrast, MTL neurons showed no significant Pre-to-Post generalization, with decoding remaining at chance level (**Figure 4f**).

### Generalization of neuronal responses for Grayscale to learned Mooney images is delayed in OC and MTL

We further analyzed the temporal dynamics of the cross-condition generalization matrices (**Figure 4a-f**). Entries along the diagonal reflect generalization from Grayscale images to Post-learning Mooney images at matched time points, while entries above or below the diagonal indicate cross-time generalization.

In the OC, the Gray-to-Post generalization matrix exhibited an asymmetric profile with a prominent off-diagonal peak (**Figure 4a**). We quantified this asymmetry by estimating the probability density function of peak decoding times via resampling (Methods). The decoding peak deviated 125 ms from the diagonal (*P* < 0.05, empirical distribution versus the diagonal; **Figure 4g**), indicating that neuronal patterns in response to Grayscale images at 175 ms most closely resembled those for Post-learning Mooney images at 300 ms (**Supplementary Figure 6a**). To evaluate whether generalization favored early or late neuronal responses, we compared decoding accuracies above and below the diagonal. Decoding accuracy was significantly higher at later compared to earlier time points in the Gray-to-Post case (**Supplementary Figure 7a,d**). By contrast, the generalization profiles for Gray-to-Pre and Pre-to-Post were centered on the diagonal and no difference was observed between early and late decoding stages (**Figure 4h,i, Supplementary Figure 6b,c, Supplementary Figure 7b,c,e,f**).

As in the occipital region, the Gray-to-Post generalization profile in MTL was also asymmetric, with higher accuracies falling below the diagonal and the decoding peak was 100 ms off-diagonal (**Figure 4j**). Thus, neuronal patterns in response to Grayscale images at 325 ms were most similar to those for learned Mooney images at 425 ms (**Supplementary Figure 6d**), and generalization was again higher at later stages (**Supplementary Figure 8**). Gray-to-Pre and Pre-to-Post generalization profiles in the MTL showed no significant asymmetry with respect to the diagonal (**Figure 4k,l, Supplementary Figure 6e,f**).

Together, these findings uncovered key neuronal dynamics induced by learning. Consistent with the behavioral findings, Grayscale-to-Mooney generalization emerged only after learning. The more pronounced below-diagonal decoding indicates that neuronal patterns for Grayscale images predominantly generalize to Post-learning Mooney images at a late processing stage. Thus, with additional processing time, responses to learned Mooney images evolve to resemble those elicited by Grayscale images. For Pre-to-Post Mooney generalization, where images are identical, the observed on-diagonal decoding in OC might reflect shared low-level visual representations.

Finally, we summarized the timing of Gray-to-Post generalization in OC and MTL (**Figure 4m,n**). Consistent with hierarchical processing, generalization occurred earlier in OC (175 ms and 300 ms, in Grayscale and Post-learning Mooney conditions, respectively) than in MTL (325 ms and 425 ms). The generalization delays were comparable across regions (125 ms in OC and 100 ms in MTL), suggesting that the temporal lag observed in OC may propagate to MTL, with MTL not contributing additional delays.

### Neurons in MTL respond later to learned Mooney images compared to Grayscale images

Both OC and MTL neuronal populations showed delayed generalization from Grayscale to Mooney images after learning. To determine whether this generalization delay was reflected in the timing of individual neuronal responses, we computed response latencies to Grayscale and Post-learning Mooney images in OC and MTL (**Figure 5a,b**). In OC, response times did not differ between conditions (**Figure 5a,c**). In contrast, MTL neurons responded significantly later to Post-learning Mooney images than to Grayscale images (time difference = 110 ms; *P* = 10^-3^, two-sided Wilcoxon signed-rank test; **Figure 5b,c**). These results suggest that MTL neurons fire only after earlier stages have devoted additional processing time to resolving the identity of the learned Mooney images.

**Figure 5.**
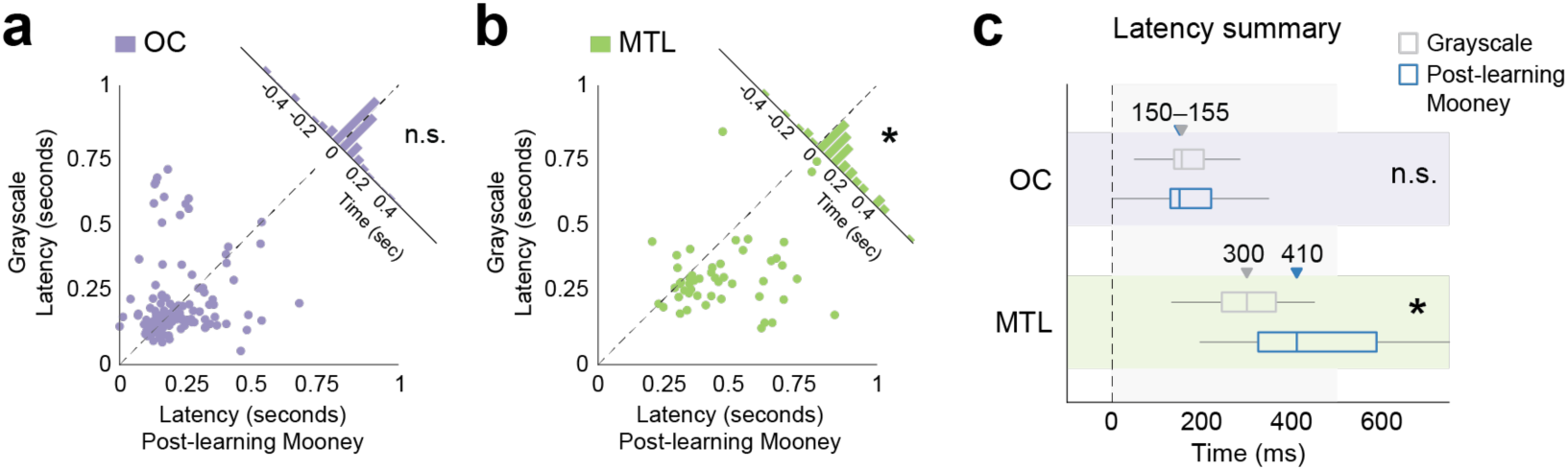
Neurons in MTL respond later to learned Mooney images compared to Grayscale images. **a**, **b**. Scatter plots of neuronal response latencies for Grayscale (y-axis) versus Post-learning Mooney images (x-axis) in the OC (**a**) and MTL (**b**). Each dot represents the response latency of a neuron to an individual image. The tilted distribution represents the differences between latencies for Grayscale and Post-learning Mooney images. The distribution centered at zero indicates no latency difference (n.s.) between conditions. The asterisk (*) in **b** denotes significant latency differences (*P* = 0.001, two-sided Wilcoxon signed-rank test) between Grayscale and Post-learning Mooney images in MTL. Thus, dots largely falling below the diagonal (dashed line) indicate that neuronal responses to Post-learning Mooney images are delayed with respect to responses to Grayscale images. **c**, Summary of neuronal response latencies. Downward arrows and accompanying numbers indicate the median latency for Grayscale (gray box) and Post-learning Mooney (blue box) images in the OC (top, purple background) and MTL (bottom, green background). In the OC, latencies did not differ between the two conditions (150 ms for Post Mooney, and 155 ms for Grayscale), and arrows overlapped. Box plots represent the interquartile range of the latency distributions in **a** and **b**. The vertical dashed line at time zero indicates image presentation onset, and the shaded gray area in the background represents the presentation period.

## DISCUSSION

We investigated how the neuronal dynamics induced by rapid learning in the human brain reshape perception using degraded two-tone Mooney images^11^. Mooney images are unrecognizable before learning. However, after brief exposure to the corresponding Grayscale images, recognition of Mooney images increases dramatically^11,12,14,16,21^. Participants in our study showed essentially all-or-none recognition of Mooney images mediated by rapid learning. Additionally, recognizing Mooney images after learning required additional processing time compared to Grayscale images.

Medial occipital neurons showed lower selectivity and faster responses, while MTL neurons showed higher selectivity and slower responses (**Figure 2**), reflecting hierarchical processing from the ventral visual pathway to downstream areas in the medial temporal lobe^27–30^. The time required for visual signals to reach the human MTL was longer compared to other primates^31–33^. Rapid learning led to changes in firing patterns in neurons in medial occipital and temporal regions. Despite the identical visual input, neurons within these areas discriminated between Pre- versus Post-learning Mooney images (**Figures 2, 3**). Previous studies in monkeys showed that neurons in inferior temporal cortex, known for their role in representing objects and faces, change their activity in response to Mooney images after exposure to their intact versions^20^, following learned associations^34–36^, and while building size-invariant object representations^37^. In humans, neuroimaging studies have revealed learning-related effects on the representation of Mooney face and object images in high-level visual cortex, including the occipital and fusiform areas, as well as in the precuneus and adjacent parietal regions implicated in top-down attention^21,24,38^. Moreover, single-shot associative memories of unrelated concepts are represented by neurons in the human medial temporal lobe^39^. Linking the work in monkeys and humans, our results demonstrate that the effects of recently acquired experience are reflected in neuronal activity in both visual cortex and MTL regions.

Following learning, neurons modulated their responses to Mooney images to resemble the responses to Grayscale images (**Figures 2, 3**). Consistently, neuroimaging studies suggested that the representation of Mooney images is more similar to the representation of intact images after learning^22,23^. Decoding Mooney images after learning required additional processing time compared to processing of intact Grayscale images (**Figure 4**), consistent with behavioral results showing slower response times for Post-learning Mooney images (**Figure 1g**). The generalization delays for degraded Mooney images (∼125 ms) are reminiscent of those previously reported in human visual areas (110-160 ms) when participants had to recognize highly occluded objects compared to whole objects^26^. Furthermore, studies in monkeys have shown that objects that are harder to recognize take longer to be reliably decoded from neurons in the primate ventral stream^40^. These delays may reflect the need for recurrent processing, involving signals from other visual areas as well as top-down feedback from areas in the frontal and temporal lobes^41–44^. Unlike these studies, our work used learning to facilitate recognition. Learning transformed an essentially impossible recognition task (Pre Mooney) into a challenging but solvable task (Post Mooney) enabling the brain to reconstruct visual objects from impoverished images. In contrast to learning new images – which may require neocortical neurons to expand their tuning to encode novel objects^45^ – recognizing Mooney images may rely on learning figure-ground segregation and image completion processes. The effects of learning in our study may manifest through extensive recurrent processing to solve object identity in the human brain, which might involve top-down feedback from higher-order cortical areas^19,46,47^.

Our analyses reveal that key neuronal dynamics that support recognition after learning occur faster in OC than in MTL highlighting their distinct roles in learning-induced visual recognition across the processing hierarchy. Pre-to-Post generalization in OC (**Figure 4**) and the early low Pre-vs-Post discriminability (**Figure 3**) may reflect shared processing of visual features of Mooney images, independent of recognition. However, once the Mooney images are learned, neuronal representations evolve to integrate the visual input with newly acquired knowledge. As a result, these dynamics lead to decoding object information in OC within 300 ms. In contrast, chance decoding observed for Pre-to-Post generalization in MTL suggests limited or no involvement in the processing of low-level visual features of the images. Instead, MTL appears to be more involved in abstract-level representations related to image identity, resolving object identity of learned Mooney images within 425 ms. Late representations after image offset (**Figure 3**) might reflect non-content-specific processing related to Mooney images after recognition^23^. Together, changes in neuronal patterns observed in OC encode rapidly learned visual information, reshaping perception through further recurrent processing before forwarding the signal to downstream areas. This process may account for the delayed neuronal responses in MTL.

Experience-driven synaptic changes in neuronal circuits are fundamental to learning^48^ and engage multiple processes and brain regions^49^. Rapidly storing memory traces is often attributed to MTL structures^18,28,39,50^, which facilitate memory reinstatement in neocortical regions via feedback^18,34^. However, whether all forms of rapid learning require the participation of the MTL remains unclear. A recent study^14^ showed that memory-impaired patients with lesions in the MTL showed performance similar to healthy control participants on a rapid one-trial perceptual learning task, suggesting cortical changes may occur independently of the MTL. Although the perceptual effects persisted for several days, the patients’ memory of the test was impaired, arguing against the involvement of associative memory^13,14^. The spatiotemporal dynamics presented here indicate that recognition after rapid perceptual learning may rely on neocortical processing without requiring feedback from MTL.

The last decade has seen remarkable progress in the development of successful Artificial Intelligence (AI) algorithms. Such algorithms are typically trained with very large amounts of data, often in a supervised fashion. Humans can use few-shot and unsupervised cues to rapidly learn, as strikingly demonstrated by recognition of impoverished two-tone Mooney images. Our single-neuron data characterize how learning-induced spatiotemporal dynamics encode new visual information reshaping perception. Unlike other forms of fast learning that require recruiting the MTL, rapid perceptual changes may primarily rely on neocortical processing. The current study provides initial steps towards understanding the dynamic interplay between visual cortex and medial temporal lobe memory structures to instantiate rapid and unsupervised learning and provide constraints for developing computational models that include recurrent computations for rapid learning.

## MATERIALS AND METHODS

### Participants

The participants undergoing neuronal recordings were 13 adult patients (34 sessions; 5 female; aged 25 to 50 years; Supplementary Table 1) with epilepsy who were implanted with hybrid macro- and microelectrodes for epilepsy monitoring and evaluation for treatment of drug-resistant epilepsy, at the Freiburg Epilepsy Center in Freiburg im Breisgau, Germany. Electrode placement was solely determined by the need to localize epileptogenic regions. The electrode locations were determined using post-implantation computed tomography co-registered with preoperative MRI. These locations were then transformed into standardized MNI152 space and plotted on a template brain for visualization purposes only (**Figure 1h**). All participants volunteered for this study and gave informed consent. The study adhered to the ethical guidelines established by the University Hospital Freiburg’s ethics committee in Freiburg im Breisgau, Germany.

### Experimental task

Participants performed an image recognition task (**Figure 1**). They were shown sequences of images and instructed to report whether they recognized the identity of the depicted objects with a button press. When recognized, participants were asked to verbally report the identity of the image. Responses were marked as incorrect if participants did not identify the image content. The images included animals, objects, and people, presented as grayscale pictures (**Figure 1b**) and their corresponding two-tone Mooney counterpart version (**Figure 1a**)^11^. At the beginning of each session, participants received instructions and were shown an example of a grayscale and the corresponding Mooney picture side by side to illustrate the images in the task. Images were presented in blocks of 60 trials, and participants completed on average 16.6 ± 3.3 blocks (mean ± s.d.) per session (**Supplementary Table 1** shows the number of sessions for each participant). Sessions stopped when a maximum of 25 blocks were reached, or earlier if a participant wanted to stop. In each trial, a grayscale or Mooney image (5° × 5° visual angle) was presented at the center of the screen for 500 ms followed by a blank screen for 1,300 ms (**Figure 1c**).

Participants were first exposed to the Mooney versions of the images (Pre-learning Mooney images, **Figure 1d**, left, red frame), which were largely unrecognized (**Figure 1e-f**). Different Mooney images were randomly interleaved over the sequence. Each Pre-learning Mooney image was presented for 30 trials before a corresponding learning segment was shown. If participants recognized the identity of a Mooney image prior to learning, that image was no longer presented, and the learning sequence for that image was not presented either. After Pre-learning Mooney images, participants were presented with a learning sequence in which an original grayscale image followed by the corresponding Mooney version of that image were used to facilitate learning (**Figure 1d**, middle, turquoise frame). If the participant recognized both the grayscale and Mooney versions of the image, that image was marked as learned. Otherwise, the learning segment was presented multiple times until both versions of the image were recognized. Once an image was marked as learned, no additional learning segments for that image were presented. For images that were recognized during the learning sequence, Mooney images were shown again multiple times (Post-learning Mooney images, **Figure 1d**, right, blue frame). Additionally, grayscale images were randomly interleaved with the Mooney images over the task. Different images from the three experimental conditions (Pre-learning Mooney, Grayscale, and Post-learning Mooney) were strategically distributed across the session to balance the number of trials per image and condition. Thus, each block within a session included an interleaved combination of all three conditions. The number of distinct images presented in each session varied depending on the number of blocks completed by participants, ranging from 4 to 10. Participants who completed multiple sessions were presented with different images in each session. Participant performance was measured as the proportion of recognized images per condition (**Figure 1e,f**). We also measured the response times as the time between stimulus onset and the participant’s button press indicating recognition.

### Eye tracking

To assess eye movements during visual recognition, we conducted the image recognition task described above in an independent cohort of healthy participants (13 sessions across 10 individuals; aged 24 to 65 years). Eye position was recorded using an infrared-based eye tracker (EyeLink 1000, SR Research) sampling at 500 Hz. Calibration was performed at the beginning of each session using a standard nine-point grid and was accepted only when the validation error was less than 1° of visual angle (mean validation error: 0.35°). The Eyelink system’s default settings were used to define fixation and saccade onsets. Visual stimuli and trial structure matched those used in the patient recordings.

### Neurophysiological recordings and spike sorting

We conducted extracellular electrophysiological recordings in patients using Behnke-Fried electrodes (Ad-Tech, Oak Creek, Wisconsin), which contained 8 microwires (40 μm in diameter) located at the tip of the electrode shaft^51–53^. Microelectrode coverage included the occipital cortex, parahippocampal cortex, entorhinal cortex, hippocampus and amygdala (Supplementary Table 2). We recorded broadband signals (0.1-9,000 Hz) at a sampling rate of 30 kHz using the NeuroPort system (Blackrock). Subsequently, these signals were filtered offline between 300-3,000 Hz using a zero-phase digital filter. We performed spike detection and sorting for each microwire using the semiautomated template-matching OSort algorithm^54^.

### Single neuron analysis

To identify visually responsive neurons, we compared firing rates during the interval from 200 to 700 ms after image onset with the baseline period, defined as the −500 to 0 ms interval before image onset. Neurons were considered visually responsive if their firing rates significantly changed in response to at least one image (either Grayscale or Mooney). Significance was assessed using a permutation test (1,000 iterations) with a P-value < 0.05, corrected for false discovery rate (FDR). FDR correction involved shuffling the labels of firing rates and repeating the analysis 1,000 times to generate a null distribution of results. The P-value threshold for visual responsiveness was then adjusted to ensure that fewer than 5% of neurons were expected to be identified as responsive by chance. Neurons in the hippocampus, amygdala, and entorhinal cortex were considered together as the medial temporal lobe (MTL). Subsequent analyses were based only on the visually responsive neurons (**Figure 2a**).

We measured neuronal response latency as the time from stimulus onset to the first significant change in firing rate relative to the baseline period (**Figure 2b**). For each responsive neuron, we estimated the instantaneous firing rate by convolving its spike train with an asymmetric kernel function ^55^ and averaging across trials. Latency was determined as the time point where instantaneous firing rate exceeded 3 s.d. from the baseline for at least 100 ms. Latency distributions were compared across regions using the Kruskal–Wallis test. Response selectivity for each neuron was defined as the number of distinct grayscale images that elicited a significant response (**Figure 2c**).

### Decoding analysis

Single-trial decoding analyses were conducted using neuronal responses as features either at the individual level (**Figure 3a,d**), or as a pseudopopulation (**Figure 3b,c,f,g**). Pseudopopulations were constructed by aggregating neurons recorded across sessions^25,26,56^. For each neuron, firing rates were first normalized by subtracting the baseline on a trial-by-trial basis and then z-scored. We used a support vector machine (SVM) with a linear kernel, as implemented by the *fitcecoc* function in MATLAB, performing five-fold cross-validation to estimate decoding performance. Decoding across time was assessed using a 250 ms sliding window, with steps of 25 ms. We built binary classifiers (chance = 50%) to discriminate between Pre-versus Post-learning conditions (**Figure 3a-c**), Grayscale versus Pre-learning (Figure 3d, f, g), and Grayscale versus Post-learning (**Figure 3e, f, g**). Each classifier label included different images. For instance, the Pre-learning label contained only Pre-learning Mooney images, but these depicted different objects, such as a camel, a tool, or people. Because different sessions contained varying numbers of images, we balanced the number of trials across sessions by randomly subsampling those with more images. We determined significance by comparing decoding performance to an empirical null distribution. The null distribution was generated by shuffling the conditions labels and running decoding for 500 iterations. Time points where decoding exceeded the 95^th^ percentile of the null distribution for at least 250 ms were considered significantly different from chance.

### Generalization analysis

We assessed generalization following a cross-condition population decoding approach, training decoders on one condition and testing them on another condition^57,58^. Support vector machines with a linear kernel were employed using a 250 ms sliding window with 25 ms steps. Pseudopopulations of neurons, pooled across sessions, were used to build classifiers. Classifiers were first trained to predict image identity from neuronal patterns in one condition. We then froze the classifier weights and tested predictions on neuronal patterns from a second condition. To account for differences in the number of images used across sessions, we chose three images from each session to build three-class decoders (chance = 33%). For each session, we sorted images based on how many neurons responded to them in the Grayscale condition and selected the top three images. To pool images across sessions, we arbitrarily labeled images in each session as Image 1-3, and then grouped those with the same labels. This approach is suitable because the SVM algorithm requires that each class elicits distinct neuronal responses to separate patterns in a high-dimensional space, independent of image identity^59^. Using this procedure, we computed cross-time generalization profiles for Gray-to-Post (**Figure 4a,d**), Gray-to-Pre (**Figure 4b,e**), Pre-to-Post (**Figure 4c,f**). For each case, we computed both possible train-test combinations. For instance, in the Gray-to-Post generalization, we first trained on the Grayscale condition and tested on the Post-learning Mooney condition to estimate decoding accuracy at each time point. Then, we repeated this process in reverse order, training on the Post-learning Mooney condition and testing on the Grayscale condition. Each entry at time *x*, *y* in the generalization matrix (e.g., Figure 4a) represents the mean decoding of these two train-test combinations. Significance was assessed by comparing decoding performance to an empirical null distribution generated by shuffling conditions labels (200 iterations). Significant decoding was defined at *P* < 0.05 (cluster-size correction for multiple comparisons)^59^.

We quantified deviation of the generalization profiles from the diagonal by computing the probability density function of peak decoding (**Supplementary Figure 6**). For each generalization profile, we resampled our data 200 times and ran our decoder as described above. Each iteration produced an x, y coordinate for peak decoding. Thus, each entry in the probability density function represents the frequency of that coordinate being identified as the maximum peak decoding. We then calculated the deviation from the diagonal (in ms) as the difference between x and y. We compared the distribution of deviations from the diagonal against the null hypothesis that these deviations are zero (**Figure 4g-l**). If 95% of the values in the distribution are greater (or lower) than zero, the deviation is considered significant. A positive deviation indicated that the patterns for the condition on the y-axis generalized to later times for the condition on the x-axis. A negative deviation indicated that the patterns for the y-axis condition generalized to earlier times for the x-axis condition. We then statistically compared the mean decoding accuracies below and above the diagonal. For each generalization profile, we selected one cluster from each side of the diagonal and compared their mean decoding (**Supplementary Figure 7, Supplementary Figure 8**). These clusters included all significant time points on both sides of the diagonal and their corresponding symmetric points. This process was repeated 200 times by resampling our data. We then employed a permutation test to compare the two distributions, determining significance at *P* < 0.05.

## ACKNOWLEDGMENTS

This work was supported by NIH grant R01EY026025, by NSF grant CCF-1231216. M.A. is supported by a postdoctoral fellowship of the Research Foundation Flanders (FWO 1299924N).

---

**Supplementary Figure 1.**
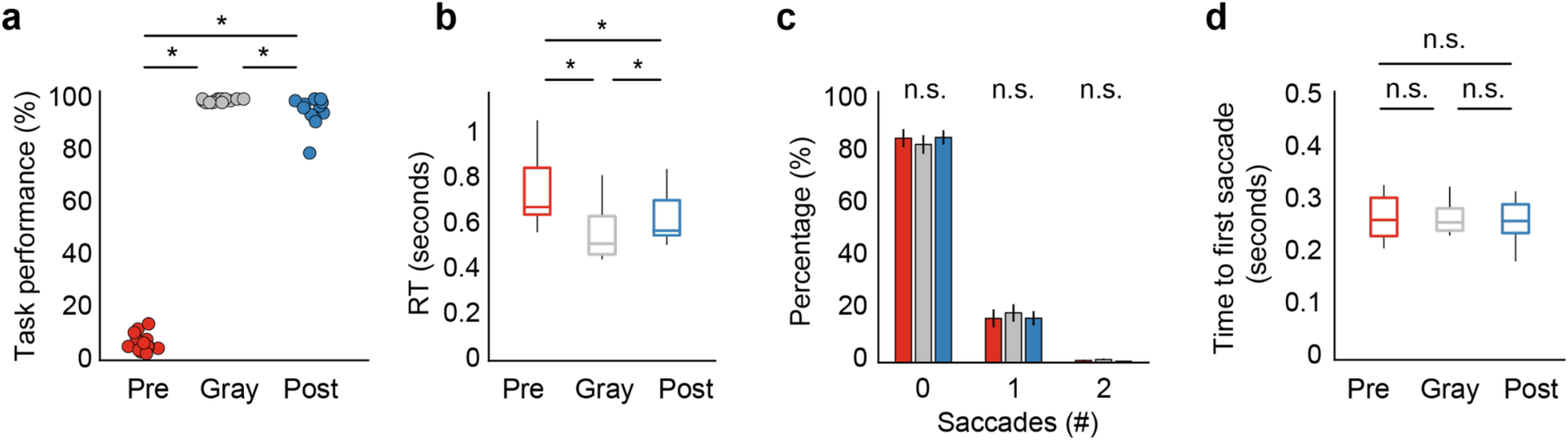
Eye movements were similar across conditions. **a**, Performance during the recognition task with eye tracking. Participants exhibited poor image recognition for Mooney images before learning (Pre, red; 6%) and nearly perfect recognition for Grayscale images (Gray, gray; 99%). After learning, recognition for Mooney images improved to 95% (Post, blue). Each dot represents one session (N = 13, 10 healthy participants, compare to **Figure 1f**). **b**, Response times (RT) relative to image onset. Participants were slower to recognize Mooney images compared to Grayscale images. Asterisks in **a** and **b** indicate statistically significant differences between conditions (*P* < 0.05, two-sided Wilcoxon signed-rank test, compare to **Figure 1g**). **c**, Percentage of trials with saccades during the image presentation interval (0–500 ms). Participants made no saccades in most trials (∼83%) and the rates of saccades were comparable across conditions (*P* > 0.05, two-sided Wilcoxon signed-rank test). Error bars are s.e.m. **d**, Latency to first saccade from image onset. When saccades occurred, they happened on average at 267 ms, and there were no significant differences across conditions (*P* > 0.05, two-sided Wilcoxon signed-rank test). Box plots show the median and interquartile range of the distribution.

**Supplementary Figure 2.**
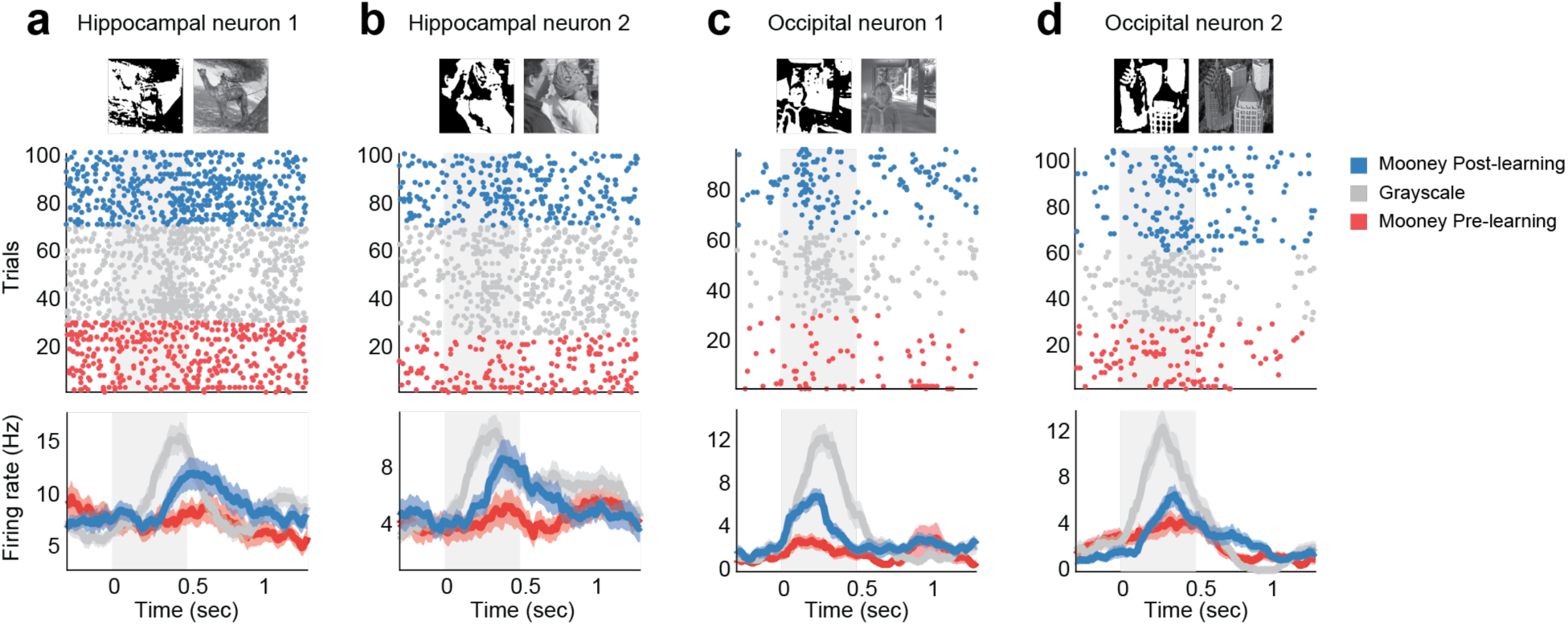
Examples of visually responsive neurons. Spiking activity of 4 example neurons in response to individual images. Raster plots (top) show spiking activity (each dot represents a spike) for an image across trials (rows) and over time. Colors correspond to spiking activity of an image across the three different conditions: Pre-learning Mooney (red), Grayscale (gray), and Post-learning Mooney (blue). The corresponding Mooney and Grayscale images are shown above the raster plots. Trials are grouped by condition for visualization. The light gray rectangle indicates the image presentation interval. Mean firing rates (bottom) are shown separately for each condition (solid colored lines; bin size is 250 ms and step size is 10 ms). Shaded areas represent s.e.m. across trials.

**Supplementary Figure 3.**
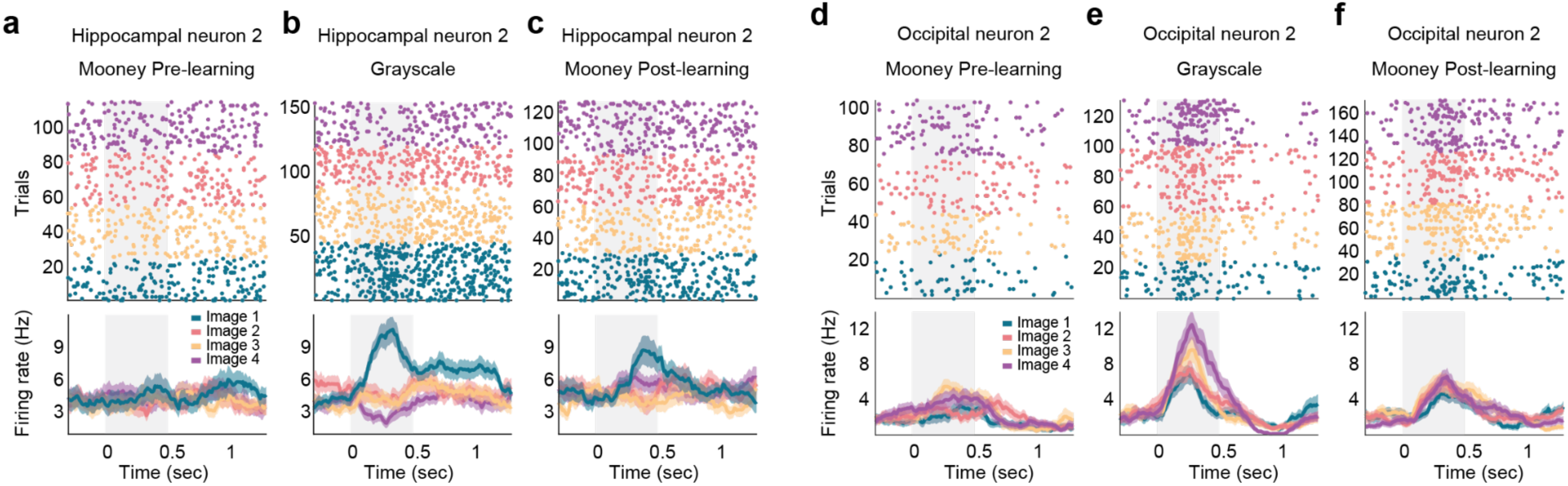
Spiking activity for example neurons across images and conditions. Spiking activity of an example neuron in the hippocampus (**a**,**b**,**c**) in response to different images in the Pre-learning Mooney (**a**), Grayscale (**b**), and Post-learning Mooney (**c**) conditions. Spiking activity of an example neuron in the occipital cortex (**d**,**e**,**f**). Data use the same conventions as **Figure 2** in the main text.

**Supplementary Figure 4.**
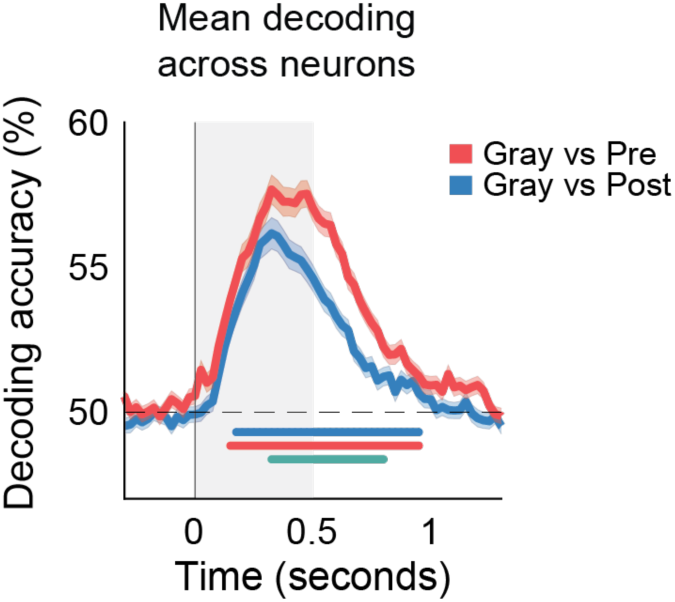
Single-neuron decoding is higher for Gray vs. Pre than for Gray vs. Post. Decoding accuracy (mean ± sem) across all neurons (n = 318) in **Figure 3d,e** for Grayscale versus Pre-learning Mooney image decoding (red) and for Grayscale versus Post-learning Mooney image decoding (blue). The dashed line is chance. Time zero indicates image presentation onset and the shaded gray area represents the presentation period. The solid lines below the curves represent time points significantly above chance for each of the curves (*P* < 0.05, permutation test). The solid turquoise line below the curves represents the time points with significant difference between the two curves (*P* < 0.05, permutation test).

**Supplementary Figure 5.**
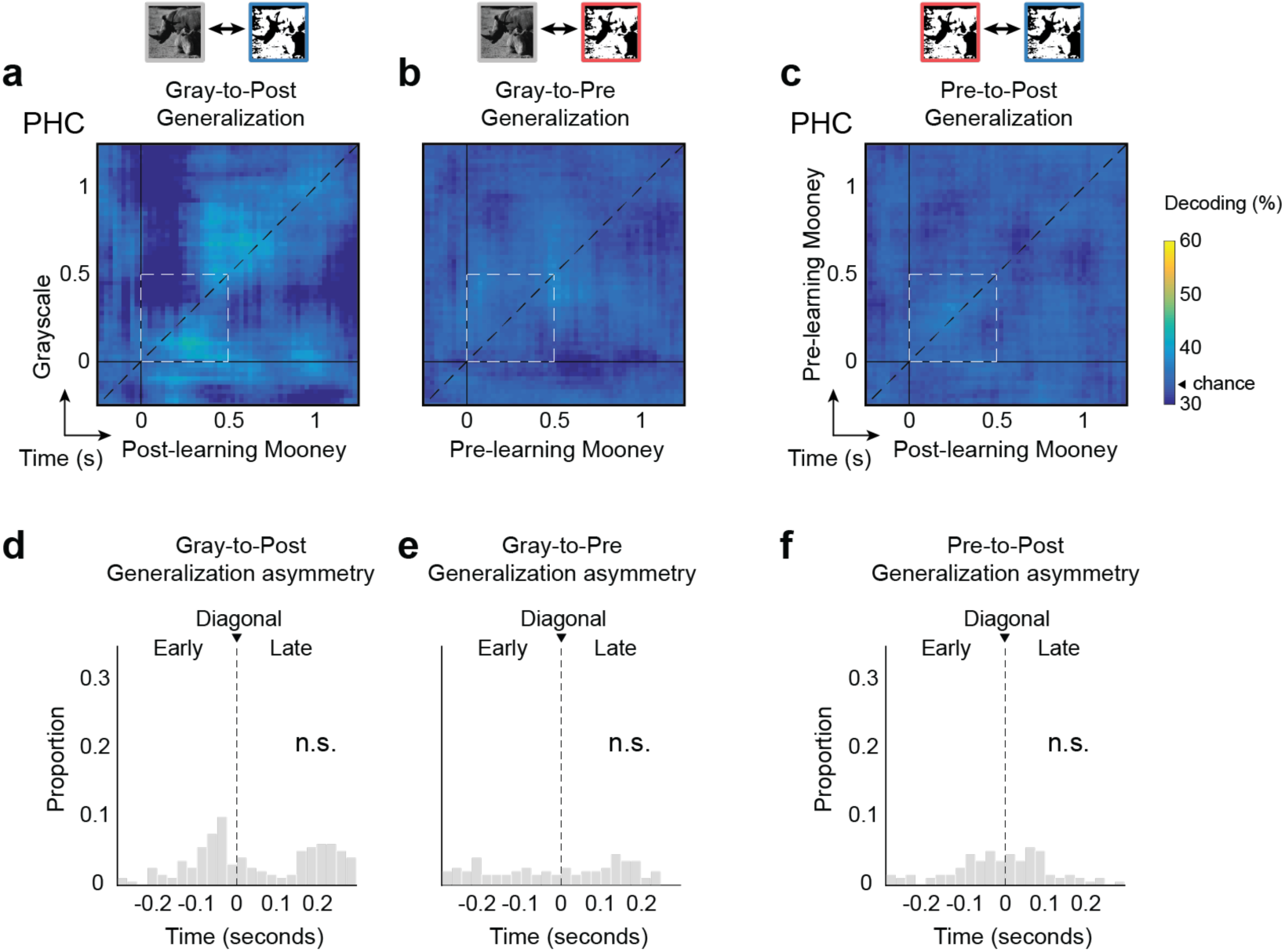
Generalization in PHC. Generalization profiles for Gray-to-Post (**a**), Gray-to-Pre (**b**), and Pre-to-Post (**c**) in PHC. Data use the same conventions as **Figure 4a-c. d**,**e**,**f**, Generalization asymmetry corresponding to the decoding profiles in **a**, **b**, and **c**, respectively. Data use the same conventions as in **Figure 4g-i**. n.s. indicates no significant deviation from the diagonal.

**Supplementary Figure 6.**
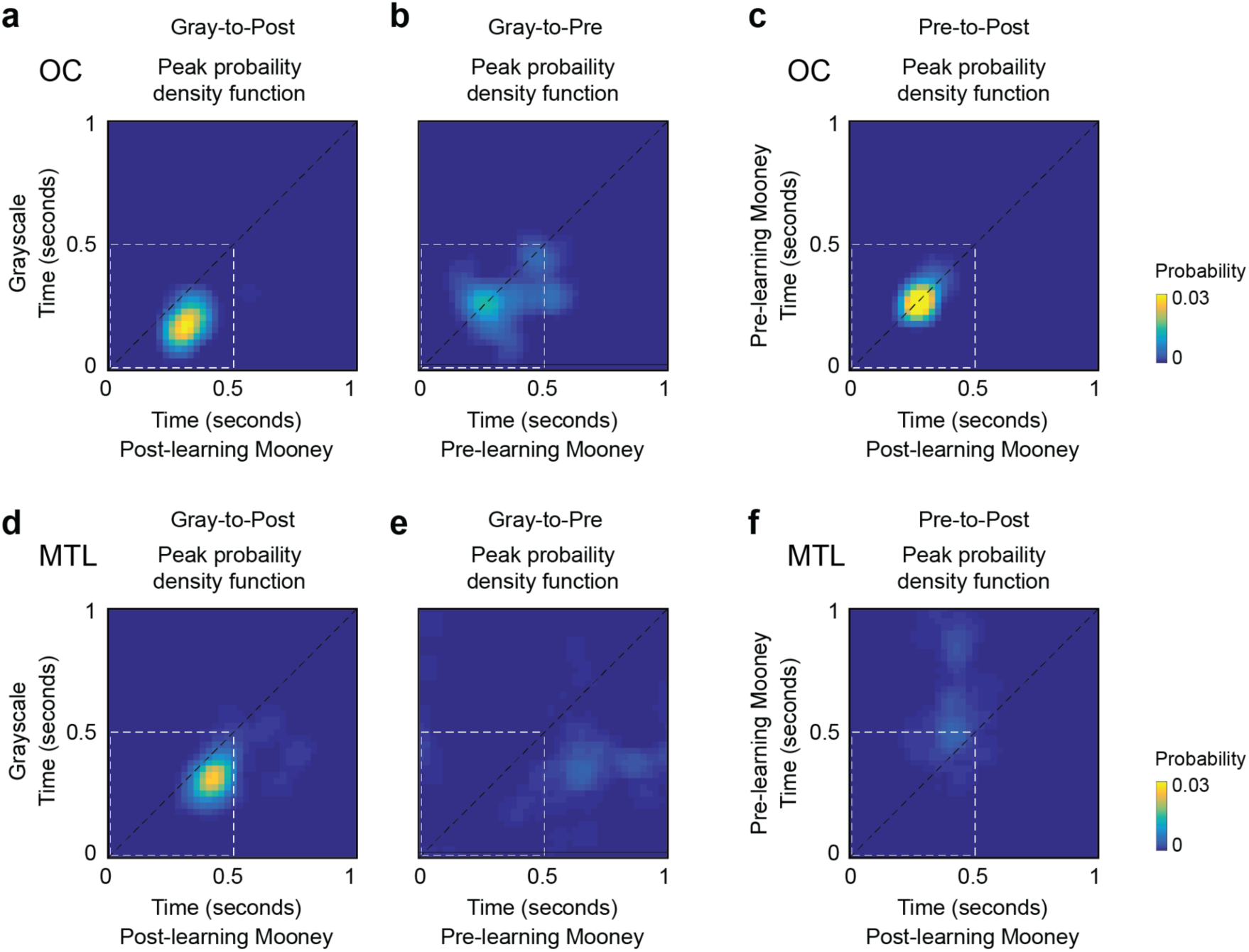
Peak decoding probability reveals asymmetric generalization dynamics in OC and MTL. **a-c**, Probability density functions of peak decoding in OC corresponding to the generalization profiles shown in **Figure 4a-c**. The matrices were computed using a resampling method to identify the maximum decoding (Methods). Each matrix entry represents the probability of observing the decoding maximum at that position. The sum of all entries in each matrix equals 1. Peak decoding below the diagonal suggests delayed generalization from Grayscale to Post-learning Mooney images. Time zero indicates image onset, the black dashed line represents the matrix diagonal, and the square with white dashed lines represents the image presentation interval.For Gray-to-Post generalization in OC (**a**), peak decoding probability is highly concentrated below the diagonal around coordinates [175, 300] ms (Grayscale and Post-learning Mooney coordinates, respectively), suggesting delayed generalization from Grayscale to Post-learning Mooney images in OC. Conversely, for Pre-to-Post (**c**), peak decoding probability falls largely on the diagonal. **d-f**, Probability density functions of peak decoding in MTL corresponding to the generalization profiles shown in **Figure 4d-f**. For Gray-to-Post generalization in MTL (**d**), peak decoding probability is highly concentrated below the diagonal around coordinates [325, 425] ms (Grayscale and Post-learning Mooney coordinates, respectively), indicating delayed generalization in MTL.

**Supplementary Figure 7.**
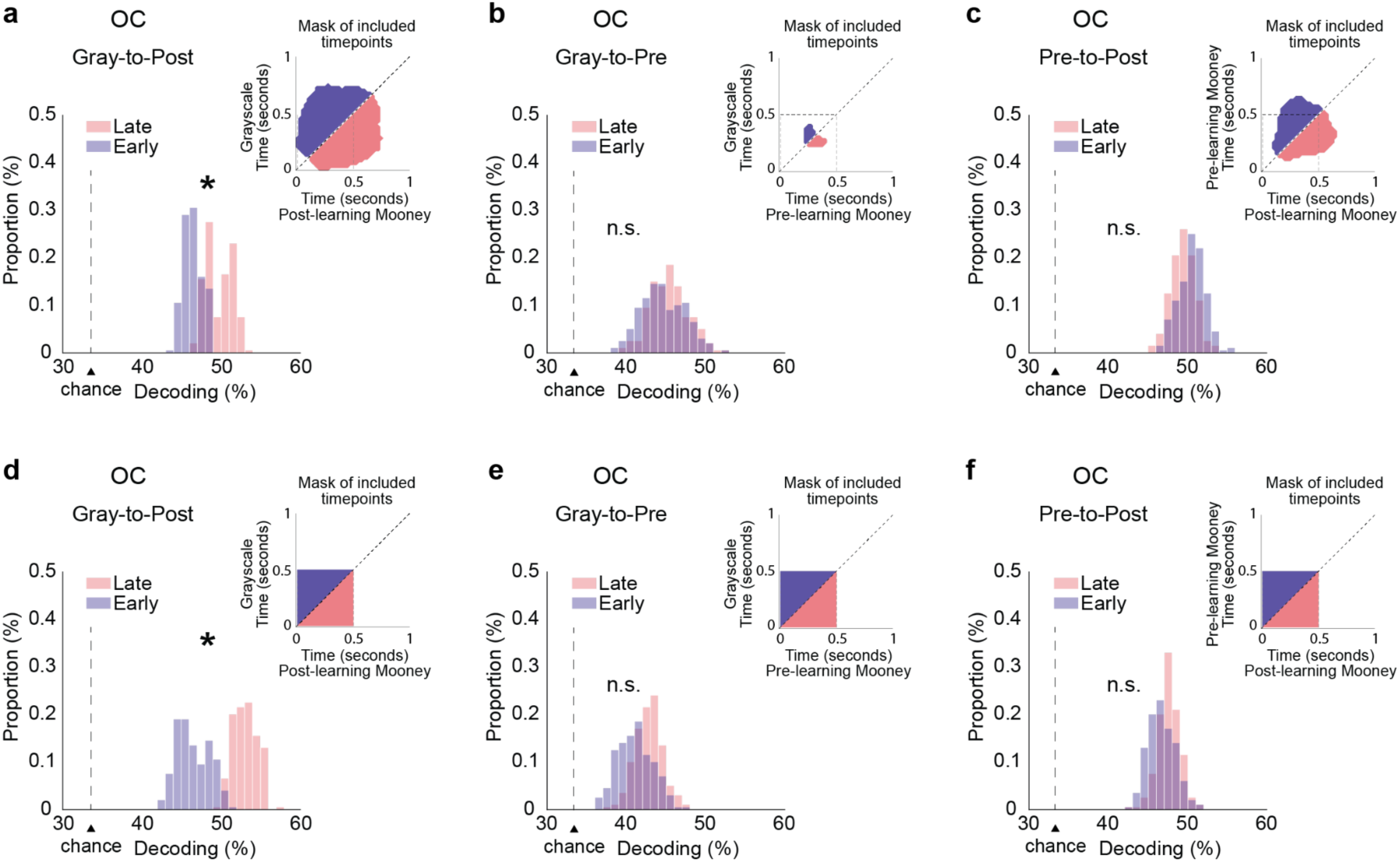
Generalization in OC is higher at late than early time points. Mean decoding accuracy below (Late) and above (Early) the diagonal for the generalization profiles in OC corresponding to matrices in **Figure 4a-c**. **a-c,** For each matrix, we selected one cluster from each side of the diagonal in the generalization matrix and compared their mean decoding. These clusters included all significant time points on both sides of the diagonal and their corresponding symmetric points. Insets show the two clusters used to compute the distributions for late (pink) and early (purple) time points. The x- and y-axis in the inset match those in the generalization matrices in **Figure 4a-c** (showing only the time range 0-1s). Distributions were computed via resampling methods. Asterisks (*) denote significant differences (*P* < 0.05) between early and late clusters. n.s. indicates no significant difference. The vertical dashed line marks decoding chance. **d-f,** Same analyses as in **a-c**, but using all time points during image presentation below (pink) and above (purple) the diagonal, as indicated in the insets.

**Supplementary Figure 8.**
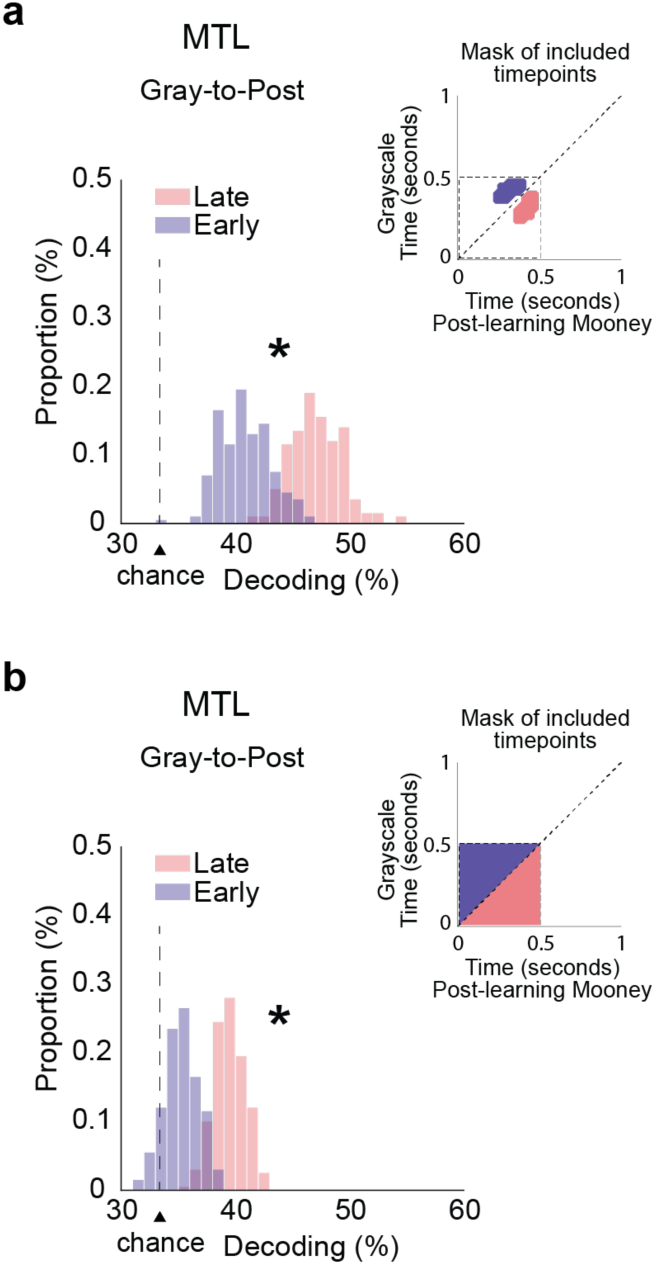
Generalization in MTL is higher at late than early time points. **a,** Mean decoding accuracy below (Late) and above (Early) the diagonal for the Gray-to-Post generalization profile in MTL corresponding to **Figure 4d. b**, Same analyses as in **a**, but using all time points during image presentation below (pink) and above (purple) the diagonal, as indicated in the insets. Format and conventions as in **Supplementary Figure 7**. A comparison was not conducted for the Gray-to-Pre and Pre-to-Post profiles in **Figure 4e,f**, given the absence of significant generalization.

**Supplementary Table 1.** Information about each of the participants in this study.

| Patient ID | Gender | Age | Handedness | # Sessions |
| --- | --- | --- | --- | --- |
| S1 | male | 25 | right | 4 |
| S2 | female | 26 | right | 4 |
| S3 | male | 50 | right | 1 |
| S4 | male | 33 | right | 1 |
| S5 | female | 41 | right | 2 |
| S6 | female | 40 | right | 3 |
| S7 | male | 35 | right | 4 |
| S8 | female | 35 | left | 3 |
| S9 | male | 48 | right | 4 |
| S10 | male | 26 | right | 2 |
| S11 | male | 45 | left | 2 |
| S12 | male | 46 | right | 1 |
| S13 | female | 33 | right | 3 |

**Supplementary Table 2.** Number of neurons recorded in each region and in each participant. OC = occipital cortex, PHC = parahippocampal cortex, Hipp = hippocampus, Amyg = amygdala, Ento = Entorhinal cortex, MTL = medial temporal lobe.

| Patient ID | OC | PHC | MTL |  |  | Total |
| --- | --- | --- | --- | --- | --- | --- |
|  |  |  | Hipp. | Amyg. | Ento. |  |
| S1 | 42 | 56 | 56 | 0 | 0 | 154 |
| S2 | 0 | 0 | 2 | 82 | 49 | 133 |
| S3 | 0 | 10 | 0 | 0 | 0 | 10 |
| S4 | 0 | 19 | 14 | 0 | 0 | 33 |
| S5 | 38 | 0 | 18 | 18 | 0 | 74 |
| S6 | 0 | 0 | 61 | 43 | 0 | 104 |
| S7 | 0 | 0 | 49 | 61 | 0 | 110 |
| S8 | 28 | 0 | 39 | 0 | 0 | 67 |
| S9 | 0 | 75 | 46 | 0 | 0 | 121 |
| S10 | 0 | 0 | 37 | 0 | 0 | 37 |
| S11 | 0 | 52 | 0 | 43 | 0 | 95 |
| S12 | 0 | 0 | 0 | 22 | 0 | 22 |
| S13 | 55 | 0 | 21 | 68 | 0 | 144 |
|  |  |  | 343 | 337 | 49 |  |
| **Total** | **163** | **212** |  | **729** |  | **1,104** |

